# Larger diet particle sizes cause crickets to grow faster with no effect on final body size

**DOI:** 10.1101/2024.08.13.607817

**Authors:** MJ Muzzatti, JD Kong, ER McColville, H Brzezinski, CC Stabile, HA MacMillan, SM Bertram

## Abstract

Artificial diets are costly to produce, so diet efficiency is critically important to the success of mass rearing insects. One way to improve feed efficiency is through dietary particle size optimization. We used a commercially reared species, *Gryllodes sigillatus*, to test whether individual crickets reared from hatch to adulthood on diets of different particle sizes would grow differently. Crickets fed a diet ≥0.5 mm grew heavier during the first three weeks but weighed the same after six weeks regardless of diet size. We then provided crickets with a choice of particle size throughout development to test for dietary size preference. Given a choice, crickets consumed the most food from the 1.0-1.4 mm diet. Crickets also preferentially select ingredients from mixed diets, so to test whether grinding a conventional diet to a finer particle size could influence performance traits, we ran a large-scale group rearing experiment and found no effect of further grinding on colony mass gain or development time. Pelleting diet is another method for eliminating self-selection of ingredients, and so we tested whether pelleting finely ground conventional cricket feed would result in any substantial changes to the developmental life history of individual crickets. Crickets fed a 2 mm pelleted diet grew larger body size but were not significantly heavier. Overall, our results demonstrate that particle size optimization can be leveraged to enhance cricket life history traits important to mass production, as growth was accelerated on larger particle size diets and crickets preferred to eat larger-sized diets. Researchers focusing on physical properties of insect diets should carefully consider the timing of growth and development through which diet particle size may influence feed efficiency.

## Introduction

Insect mass rearing facilities raising mealworms, crickets, and black soldier flies are relatively new to Europe and North America and are gaining traction. The biggest hurdle facing the success of the insects as food and feed industry is scaling up production to meet demand (Larouche *et al*., 2023). Successfully rearing insects that develop into healthy adults who contribute offspring to the next generation is largely dependent on the development of an artificial diet that a mass-reared species will willingly consume (Cohen, 2015; Morales-Ramos *et al*., 2014). High-quality diets are often difficult to produce in large quantities and diet remains one of the most expensive components of mass rearing facilities (Aceituno-Medina *et al*., 2020; Morales-Ramos *et al*., 2014, 2020); for example, diet in swine or poultry systems contributes to 65 – 75% of overall production costs (Goodband et al., 2002). Changes in diet can help reduce these costs (van Huis, 2022). There is great potential to optimize farmed insect production through diet, but while we know there are strong effects of diet quality on insect growth and development (Cammack and Tomberlin, 2017; Harrison *et al*., 2014; Serrato-Salas and Gendrin, 2023), far less is known about the impacts of the physical properties of diet on insects.

Modifications to diet physical properties can help ensure that carefully selected and expensive ingredients that match the nutritional profile of the insect are consumed and not wasted. Such improvements in feed efficiency can lower production costs even when diet content is unchanged (Jo *et al*., 2021). Feed efficiency in farmed animals can be improved, for example, through dietary particle size optimization. Dietary particle size is an important nutrition factor in many agricultural systems. In poultry, for example, optimum particle sizes are dependent on the feed ingredients (ex, maize, sorghum), and can influence substrate availability for enzymatic digestion (Amerah *et al*., 2007). In pigs, particle size reduction in feed results in increased feed efficiency, resulting from improved nutrient digestibility (Goodband *et al*., 2002; Jo *et al*., 2021). This is unsurprising, given a food’s surface area to volume ratio is increased when it is broken into smaller particles; because digestive enzymes have more surface area to act upon, more nutrients are potentially released from the food (Berthaume, 2016).

If modulation of particle size can improve yields of vertebrates in a farm setting, the same may be true for insects. In insects, particle size impacts diet retention time in the gut, modulating nutrient availability (Cohen, 2015; Parada and Aguilera, 2007; Tennis, 1985; Tennis *et al*., 1979). As a result, insects can vary in response to food particle size throughout development in ways that influence performance and fitness. For example, in mass reared tephretid flies (*Anastrepha obliqua*), larger particles of corn cob, but not coconut fiber, increased the number of larvae recovered per gram of diet fed and the percent adult emergence (Rincón-Betancurt *et al*., 2022). Black soldier fly larvae development is also influenced by food particle size, as they develop slower when fed substrates with smaller particle sizes (Yakti *et al*., 2023).

Food particle size can also impact insect accessibility to the diet, as insects must use their mouthparts to obtain access to the food (Cohen, 2015; Parada and Aguilera, 2007; Tennis, 1985; Tennis *et al*., 1979). As a result, insects demonstrate preferences for diet particle size. Dung beetles will select smaller dung particles compared to the maximum particle size that can be ingested, a feeding strategy thought to increase the efficiency of the filter feeding mechanisms by reducing the amount of undigested fibres that require chewing (Holter, 2000; Holter and Scholtz, 2007; Madzivhe *et al*., 2021). Ants also demonstrate species-specific preference for particle size, and this response is related to forager head width; species with wider heads prefer larger particle sizes (Hooper-Bùi *et al*., 2002). Together, these pieces of evidence suggest that dietary particle size can influence life history traits of insects in a species- and ingredient-specific manner.

Teasing apart the effects of particle size from other aspects of the diet can be complex. Consider, for example, the study by Tennis and colleagues who fed small- and large-strain populations of house crickets (*Acheta domesticus*) chicken laying mash diets. The small cricket strain had greatest fitness and grew heavier most quickly during the first stage of growth when fed the small particle diet. Conversely, the large cricket strain had greatest fitness and grew heavier most quickly during the last stage of growth when fed the large particle diet (Tennis et al. 1979). Further, across multiple generations, crickets fed the small particle diet were smaller than crickets fed the large diet. While this experiment suggests food particle size acts as a selection pressure on body size, there is another potential explanation (Tennis, 1985; Tennis *et al*., 1979). Chicken laying mash is a heterogenous mix of grains and seeds. The large particle food may have enabled crickets to preferentially consume some types of grains and seeds over others, altering the nutrient ratio of the diet. Such a preference for dietary particle size in crickets has been previously suggested (Patton, 1963), but not experimentally tested.

Different ingredients have different physical properties, such as hardness or brittleness, and so the same level of milling/grinding can result in different ingredients having different sized particles. The resulting heterogeneity of cricket diets means that self-selection behaviour could be driven by a preference for particle size, a preference for specific ingredients contained within the diet, or a combination of the two. Finely grinding and then pelleting conventional cricket feeds is one possible solution for preventing size-selective feeding in farmed insects like crickets. Pelleted diets may offer advantages such as improved nutrient distribution, a uniform and consistent diet, minimized time and labour costs associated with handling and weighing specific ingredients, and decreased microbiological contamination (Aceituno-Medina *et al*., 2020). Pelleted diets promoted weight gain of larvae and pupae of tephretid flies (*Anastrepha obliqua* and *A. ludens*), two insect species mass-reared for sterile insect technique (Aceituno-Medina *et al*., 2020). Other insects are successfully reared on pelleted artificial diets, including silkworm (*Bombyx mori*; Ohura and Li, 2001), maize weevil (*Sitophilus zeamais*; Ojo and Omoloye, 2012), swallowtail butterfly caterpillars (*Papilio xuthus*; Kim, Seong Hyeon et al., 2014), and Angoumois grain moth (*Sitotroga cerealella*; (Chippendale, 1971a; Chippendale, 1971b). It is unclear, however, whether a pelleted diet may offer advantages for cricket production.

We hypothesized that performance traits important to mass production of a commercially reared cricket species, *Gryllodes sigillatus*, could be enhanced by either altering the food particle size or by pelleting the food. Crickets are generalist omnivores, possessing biting mouthparts adapted for chewing and breaking down solid animal or plant materials to smaller ingestible pieces (Cortes Ortiz *et al*., 2016; Ogita *et al*., 2021). Typical cricket feeds contain a mix of corn, soymeal, fishmeal, vitamins, and minerals (Morales-Ramos *et al*., 2020), and each ingredient is individually milled to an ingredient-specific particle size range prior to mixing. We performed a series of experiments that tested how altering some of the physical properties of diet, such as particle size and pelleting, influences the life history traits of *G. sigillatus*. In our first experiment we controlled for heterogeneity of nutrients within a diet by rearing individual crickets from hatch to adulthood on commercial rabbit kibble that we had ground and sieved into one of three different particle sizes. We used rabbit kibble to produce these diets as it is relatively homogenous in nutrients compared to typical farm diets, which enabled us to isolate the effects of particle size from the effects of preferential nutrient selection. We predicted that crickets fed the smaller particles would grow the largest because small particles have increased surface area for enzymatic digestion compared to medium and large particles. In a follow-up to the first experiment, we tested whether crickets demonstrated a preference for particle size throughout their development by giving them an opportunity to consume all three particle size diets and quantifying how much of each was consumed. We predicted that crickets would demonstrate a dietary preference for particle size, and this preference would scale with growth; crickets would prefer smaller sized diets early in life, and larger sized diets later in life. In our second experiment, we tested whether finely grinding and then pelleting a conventional cricket feed (containing the traditional mix of corn, soymeal, fishmeal, etc.) would result in any substantial changes to developmental life history of individual crickets. We predicted that crickets fed the pellets would grow the largest because the pellet restricts selective feeding ensuring the cricket consumes all nutrients within the diet. In our third experiment, we tested whether reducing the choice for particle size selection by finely grinding a conventional cricket feed affects the growth and development of crickets. We predicted that crickets fed a finer ground diet would grow the largest because they would be restricted from leaving larger particle sizes behind.

## Materials and Methods

All experimental crickets were received as eggs from a commercial cricket supplier (Entomo Farms, Norwood, Ontario, Canada). The colony was fed ad libitum water and a proprietary commercial cricket diet made mostly from corn, soybean meal, linseed meal and fishmeal. Eggs were laid in a medium of peat moss and maintained inside an incubator (Thermo Fisher Scientific Inc., Massachusetts, United States) at 32°C, 60% relative humidity, and 14 h:10 h (light:dark) until hatching. These conditions were also maintained throughout development for all experiments except for the group-rearing fine grinding experiment which was maintained in a greenhouse at 30°C, 20% relative humidity, and 14 h:10 h (light:dark).

### Particle size experiments

Three diets different in particle size were created by grinding uniformly sized rabbit kibble (Martin Little Friends, Martin Mills Inc., Elmira, Ontario, Canada; full ingredient list and proximate analysis found in Table S1 and Table S2) (a laboratory cricket diet; Judge et al. 2008) through a household coffee grinder (Cuisinart CBM-18C Programmable Conical Burr Mill). We ground the rabbit kibble using the finest grind setting and the coarsest grind setting. We then mixed these finely- and coarsely-ground together and then passed it through a sieve series (1.4 mm, 1.0 mm, 0.7 mm, 0.5 mm, 0.125 mm, 0.088 mm; Figure S1) using a mechanical sieve shaker (Gilson 8 in Sieve Shaker, 115V/60Hz). Our three diet treatments spanned a 10-fold difference in particle size: small (0.088 – 0.125 mm), medium (0.5 – 0.7 mm), and large (1.0 – 1.4 mm). We also tested a fourth diet treatment, as we also reared crickets on the small diet for the first 28 days of their development and then reared them on the large diet for the remainder of the experiment.

We tested the impact of diet size on cricket performance by individually rearing 40 *G. sigillatus* from egg to adulthood on each diet treatment, resulting in a total of 160 crickets. Crickets were housed in 7.4 × 3.6 cm (top diameter x height) 96.1 mL clear plastic condiment cups with aerated lids (Solo Cup Company, Illinois, United States) and were provided with egg carton for shelter. Crickets were weighed using a Mettler Toledo model AB135-S analytical scale and photographed for body size measurements using a Zeiss Stemi 305 Stereo Microscope with an Axiocam 208 camera at hatch, three weeks and six weeks post-hatch. Body size measurements were performed on photographs using ImageJ, version 1.48 software (National Institutes of Health, Bethesda, Maryland, United States of America), and included head width, pronotum width, and pronotum length. Measurements were averaged across three independent measurers to reduce measurement error. Principal component analysis was used to quantify adult body size by creating a single summary variable (PC1) for all three body size measurements (Muzzatti *et al*., 2022). The first principal component explained 98.8% of the variation in size (eigenvalue = 2.96), and all size measures loaded equally on the first principal component. Crickets were monitored for death and time to adulthood thrice weekly. Upon eclosion to adulthood crickets were weighed and photographed again. Crickets were provided *ad libitum* fresh food and water on a weekly basis, and we also quantified weekly food intake (difference in weight of food dish before and after consumption). Feed dishes consisted of a 14-mm-wide polyethylene push-in cap glued to one half of a 5 mm wide petri dish for a food dish. To accurately measure the amount of food consumed, before crickets were provided with their food, we ensured it was dried in an oven containing dishes of blue indicating silica gel desiccant beads (Dry & Dry, Brea, California, USA) at 30°C for 96 h. We repeated this process with the partially eaten food again after it had been in the individual’s container for a week of consumption. Frass was manually removed from post-feeding dishes using forceps prior to weighing.

Normality for days to adulthood, body size, mass, and food consumption measurements was confirmed by inspecting the residuals, and all statistical analyses were performed in RStudio version 2023.12.1 (RStudio Team, 2020). We used a survival analysis followed by pairwise comparisons using a Log-Rank test and a Bonferroni p-value adjustment to test how diet particle size influenced the development time to adulthood.

We used a linear model at Week 0 to ensure that the nymphs selected for each diet treatment were the same mass. To test if dietary particle size influences mass and body size we used linear models, one at week 3 and one at week 6. We chose a linear model instead of a mixed model approach because cricket size and mass were only measured twice. At Week 3, crickets on the switch diet were grouped together with crickets on the small diet, because up until this point both treatment groups were treated the same and fed the small diet. At Week 6, the crickets on the switch diet were treated as a separate treatment. Both models included sex and the interaction between sex and diet as fixed effects. Pairwise comparisons for significant model effects were created using Tukey’s Honestly Significant Difference tests.

We used linear mixed-effects models using the ‘lme4’ package (Bates *et al*., 2015) to test if dietary particle size influenced amount of food consumed. We used a mixed model because the within-subject observations might be correlated across the weekly consumption data. The mixed model considers the switch diet treatment crickets separate from the small diet treatment crickets even though they were all fed the small diet from weeks 0-4. We included diet, week, sex, and interactions between the three as fixed effects, and cricket ID as a random effect. We performed model selection and identified the ‘best approximating model’ for amount of food consumed by calculating Akaike weights to determine the relative likelihood of the current model and by calculating the evidence ratio to determine how many times more parsimonious the top ranked model is over the current model (Table S6). Type III analyses of variance were fit to the best approximating model to test for overall differences of model terms. Pairwise comparisons were created using Tukey’s Honestly Significant Difference tests.

To test for preferences in particle size, we used the same three diets: small (0.088 – 0.125 mm), medium (0.5 – 0.7 mm), and large (1.0 – 1.4 mm). Upon emergence, 50 crickets were individually housed in 18.4 × 12.7 × 5.08 cm (length x width x height) 709.8 mL clear plastic take-out containers with aerated lids (Platinum Crown Corporation Limited, Colchester, United Kingdom). Crickets were provided with egg cartons for shelter and with separate feed dishes for each particle size diet. Each cricket was allowed to feed *ad libitum* across all diets throughout development. The positions of the feed dishes were maintained across all containers and were rotated weekly. Crickets were provided with fresh food weekly and water biweekly for six weeks. Weekly food consumption was measured using the procedure described above. Week one measurements were omitted from analysis as hatchlings consumed minimal food.

Crickets were photographed weekly for body size measurements and weighed at the end of the experiment following the same procedure outlined in Experiment 1A. PC1 explained 99.65% of the variation in size (eigenvalue = 2.98), and all size measures loaded equally on the first principal component. Normality of body size, mass, and food consumption measurements was confirmed by inspecting the residuals, and all statistical analyses were performed in RStudio version 2023.12.1 (RStudio Team, 2020). We used a linear mixed-effects model using the ‘lme4’ package (Bates *et al*., 2015) to test how cricket preference for dietary particle size changed over time. We used diet, week, sex, and interactions between the three as fixed effects, and included cricket ID as a random effect. We performed model selection and identified the ‘best approximating model’ using the same procedure from the first experiment (Table S7). A type III analysis of variance was fit to the best approximating model to test for overall differences of week and diet, and pairwise comparisons were created using a Tukey’s Honestly Significant Difference test. Normality was confirmed through residual analysis.

### Pellet experiment

To test the influence of pelleting feed on cricket life history traits, we created a cricket kibble. We used a commercial cricket feed (Earth’s Harvest Organic Cricket Grower; Earth’s Harvest, Oxford Mills, Ontario, Canada; full ingredient list in Table S8) and ground it into a fine powder using an electric coffee grinder. The ground diet was then mixed with tap water in a 1:1 ratio to create a paste, which was then smeared over 1 mm thick plexiglass sheet molds (Langaelex, Guangdong, China) with 3 mm diameter holes drilled throughout. The molds were placed in a 30°C drying oven for 48 h, and the resulting pellets were gently popped from their molds.

Upon emergence, 100 crickets were individually housed in 7.4 × 3.6 cm (top diameter x height) 96.1 mL clear plastic condiment cups with aerated lids (Solo Cup Company, Illinois, United States), provided with egg carton for shelter, and watered and fed *ad libitum* one of two diets: the ground or pelleted cricket feed. Crickets were weighed weekly for six weeks and photographed at the end of the experiment for body size measurements, following protocols outlined in the first experiment. PC1 explained 91.36% of the variation in size (eigenvalue = 2.74), and all size measures loaded equally on the first principal component. Normality of body size and mass measurements was confirmed by inspecting the residuals, and all statistical analyses were performed in RStudio version 2023.12.1 (RStudio Team, 2020). We used a linear mixed-effects model using the ‘lme4’ package (Bates *et al*., 2015) to test whether pelleting cricket feed influences cricket mass over time. We used diet, week, sex, and some of their interactions as fixed effects, and cricket ID as a random effect. We performed the same model selection process as mentioned previously (Table S9), and pairwise comparisons were created using a Tukey’s Honestly Significant Difference test. We also used linear models to test whether pelleting cricket feed influence cricket body size and mass at the end of the experiment (Week 5) and included sex and the interaction between sex and diet as fixed effects. Pairwise comparisons for significant model effects were created using Tukey’s Honestly Significant Difference tests.

### Fine grinding experiments

To test the effect of fine grinding cricket feed on cricket life history traits, we ground the same commercial cricket feed used in the pellet experiment using a commercial food-grade 304 stainless steel grain grinder (50 kg, 2200W; VEVOR, Shanghai, China) on the finest possible setting (200 mesh). Upon emergence, 79 crickets were housed in the exact same manner as the pellet experiment and watered and fed *ad libitum* one of two diets: the fine grind or the untreated commercial cricket feed. Crickets were weighed weekly for seven weeks, and survival and time to adulthood were monitored three days a week. A total of 62 crickets survived after seven weeks and were used in statistical analyses. We also tested the effect of fine grinding at the colony level by seeding four replicate bins (61 × 41 × 42cm) with 500 first instar crickets for each feed treatment (unground control and ground). Treatments were randomly allocated to bins and the bins were maintained in a greenhouse for seven weeks. Weekly, 20 crickets were randomly sampled from each bin, weighed, and instar number recorded. A final census was recorded after seven weeks, and the total mass (biomass) of the remaining crickets from each bin was recorded.

To analyze differences in mass, we fitted weighted logistic growth curves to mass over time with a power variance function for weighting wet mass using nlme v3.1 (Pinheiro and Bates, 2000). Two models were constructed: a model pooling across all individuals and feed type representing a null hypothesis, and a model fitted to each feed type pooling across individuals representing an alternative hypothesis. The alternative hypothesis model was tested against the null hypothesis model using a likelihood ratio test. Sex-specific mass data were analysed using analyses of variance with an interaction term between sex and instar and an additive effect of feed type. A three-way interaction between instar, sex and feed type was not significant for group-reared (F_2, 666_ = 0.16, *P* = 0.85) or individually reared crickets (F_2, 191_ = 4.84, *P* = 0.61) in an initial analysis. Development time to adulthood and survival were analysed using binomial regressions. Biomass at week seven was analyzed using a t-test for group-reared crickets only.

## Results

### Particle size experiments

Of the 40 newly hatched nymphs fed each diet treatment testing the impact of diet size on cricket performance (160 in total), 6 went missing (Small: 3, Medium: 1, Large: 1, Switch: 1), 56 died prior to adulthood (Small: 12/37, Medium: 15/39, Large: 14/39, Switch: 15/39), 8 were still juveniles by the end of the six weeks (Small: 4, Medium: 0, Large: 1, Switch: 3), and 90 developed into adults (Small: 21/37, Medium: 24/39, Large: 24/39, Switch: 21/39). A single cricket on the small diet developed into an adult but was lost before it could be weighed and photographed at adulthood, and so the total adult sample size was reduced to 89.

There were significant differences in the time to adulthood between particle size diet treatments (log-rank test: X^2^ = 9.5, df = 3, *P* = 0.02; Figure 1). Crickets fed the large diet developed into adults significantly faster than crickets fed the small diet (*P* = 0.046). The average number of days required to reach adulthood (± standard error) for crickets fed each diet were small = 35.75 ± 0.74 days, medium = 35.21 ± 0.54 days, large = 33.58 ± 0.43 days, and switch = 35.71 ± 0.63 days. Cricket mass did not differ among diet particle size treatments at the start of the experiment (F_3,90_ = 1.26; *P* = 0.30). However, by week three, diet particle size significantly influenced mass (F_2,86_ = 11.09; *P* < 0.001; Figure 2; Table S3). At this time, crickets fed the medium and large diets were 30.6% and 41.8% significantly heavier than crickets fed the small diet, respectively (Figure 2; Table 1). By week six diet particle size no longer significantly influenced mass (F_3,81_ = 1.14; *P* = 0.34; Figure 2; Table S3). There were no significant interactions between sex and diet at week three or six. Body size at week three showed a similar pattern to mass: diet particle size significantly influenced body size (F_2,84_ = 10.11; *P* < 0.001; Figure 3; Table S3); crickets fed the medium and large diets had 34.5% and 36.2% significantly larger bodies, respectively, than those fed the small diet at this time (Figure 3; Table 1). As with mass, by week six there was no longer a significant effect of diet on body size (F_3,69_ = 0.73; *P* = 0.54; Table S3). Females and males did not differ in size at week three (F_1,84_ = 0.075; *P* = 0.78), but females were significantly larger than males by week six (F_1,69_ = 50.04; *P* < 0.001); neither interaction between sex and diet were significant at either week three or six (Table S3). Diet particle size did not significantly influence the amount of food consumed (F_3,89.49_ = 0.68; *P* = 0.57; Figure 4). However, the amount of food consumed was significantly influenced by week of study providing the unsurprising realization that as nymphs age and grow they consume more food (F_5,450.51_ = 208.99; *P* < 0.001). Females ate significantly more food than males (F_5,450.51_ = 208.99; *P* < 0.001).

**Table 1.** Least squares means ± standard error of adult mass and body size (PC1) of individual crickets from the no-choice diet particle size experiment. Values within a column followed by different letters are significantly different (*P* < 0.05).

| Week | Diet | Mass (g) |  | Body size (PC1) |  |
| --- | --- | --- | --- | --- | --- |
| 3 | Small | $0.052 \pm 0.003$ | A | $-1.77 \pm 0.10$ | A |
| | Medium | $0.067 \pm 0.004$ | B | $-1.16 \pm 0.15$ | B |
| | Large | $0.076 \pm 0.004$ | B | $-1.13 \pm 0.14$ | B |
| 6 | Small | $0.25 \pm 0.017$ | A | $1.64 \pm 0.13$ | A |
| | Medium | $0.26 \pm 0.014$ | A | $1.51 \pm 0.17$ | A |
| | Large | $0.25 \pm 0.015$ | A | $1.69 \pm 0.13$ | A |
| | Switch | $0.25 \pm 0.012$ | A | $1.51 \pm 0.13$ | A |

**Figure 1.**
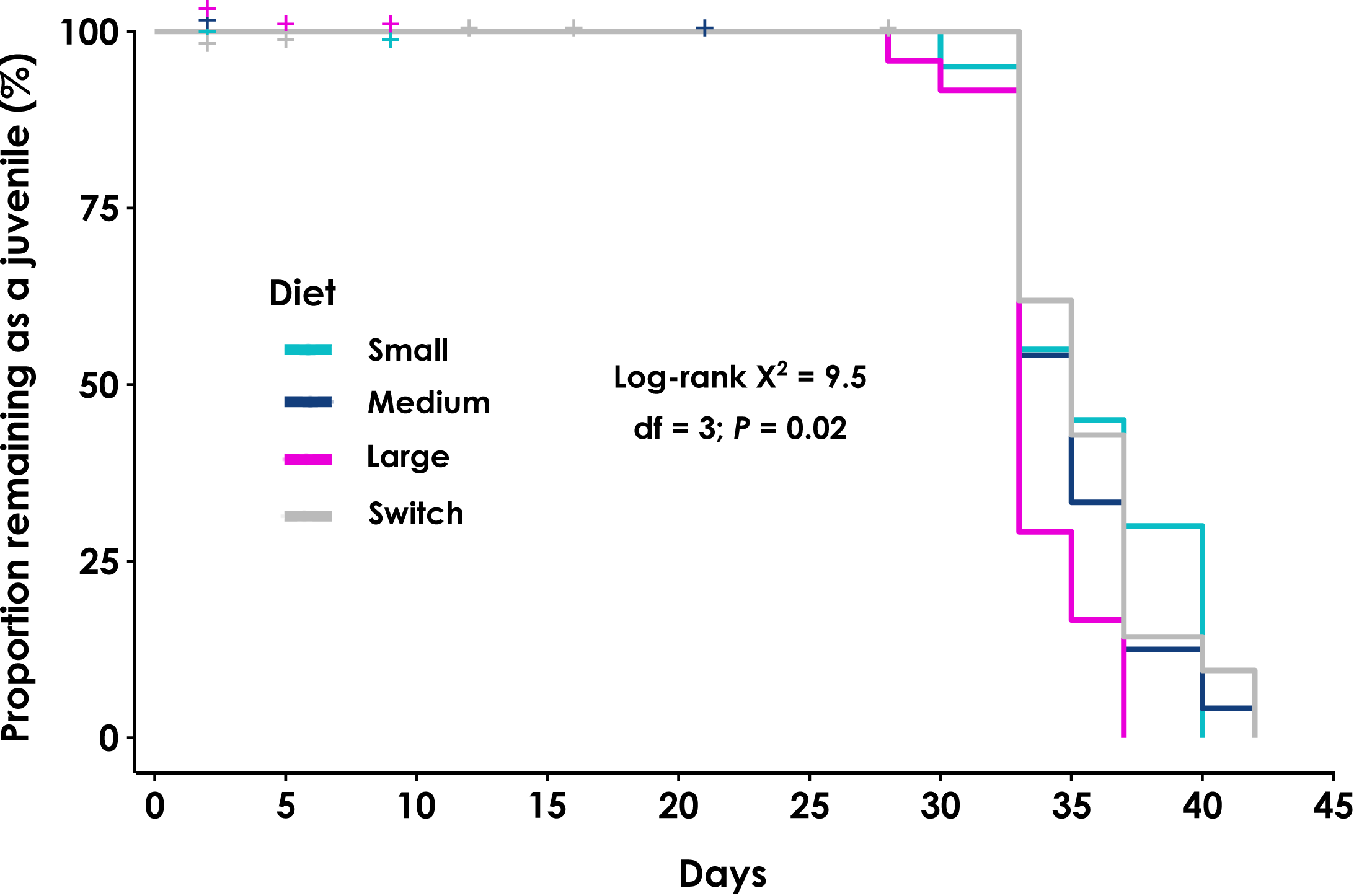
Kaplan-Meier plot representing the proportion of crickets remaining as a juvenile by a certain date. Individual crickets were fed three different sizes of rabbit kibble: small (0.088-0.125 mm), medium (0.5-0.7 mm), large (1.0-1.4 mm), and a switch from small to large at 28 days after hatch. The curves represent N=145 total crickets, with N=89 that developed into adults (Small: 20/37, Medium: 24/39, Large: 24/39, Switch: 21/39). The steps in each diet curve represent the day crickets eclosed into adults. Ticks on the curves represent censored individuals that either died or went missing before developing into adults. Jitter was applied along the y-axis to display overlapping ticks.

**Figure 2.**
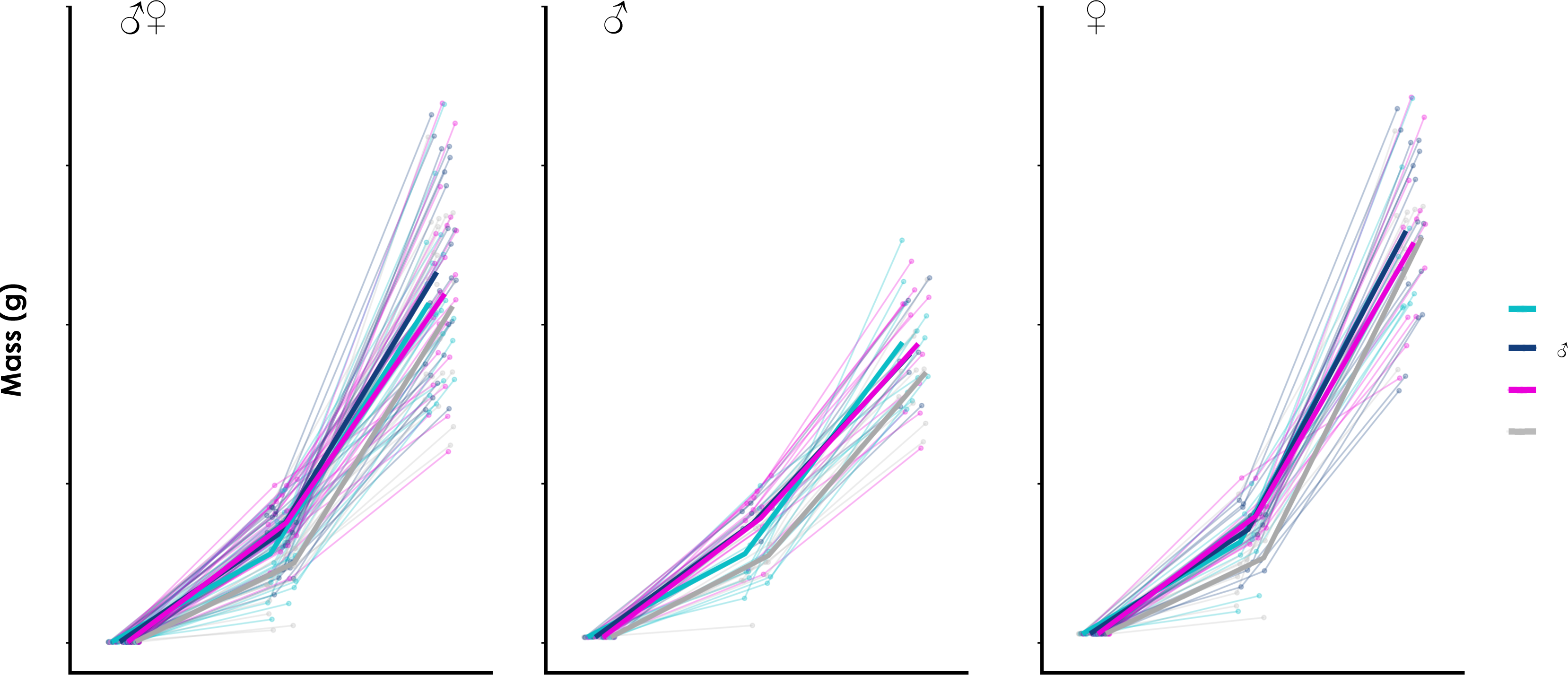
Lifetime mass as a function of age of Gryllodes sigillatus fed one of four different sizes of rabbit kibble: small (0.088-0.125 mm), medium (0.5-0.7 mm), large (1.0-1.4 mm), and a switch from small to large at 28 days after hatch. Thick lines connect mean values at each age and solid circles represent individual crickets connected by thin lines over time.

**Figure 3.**
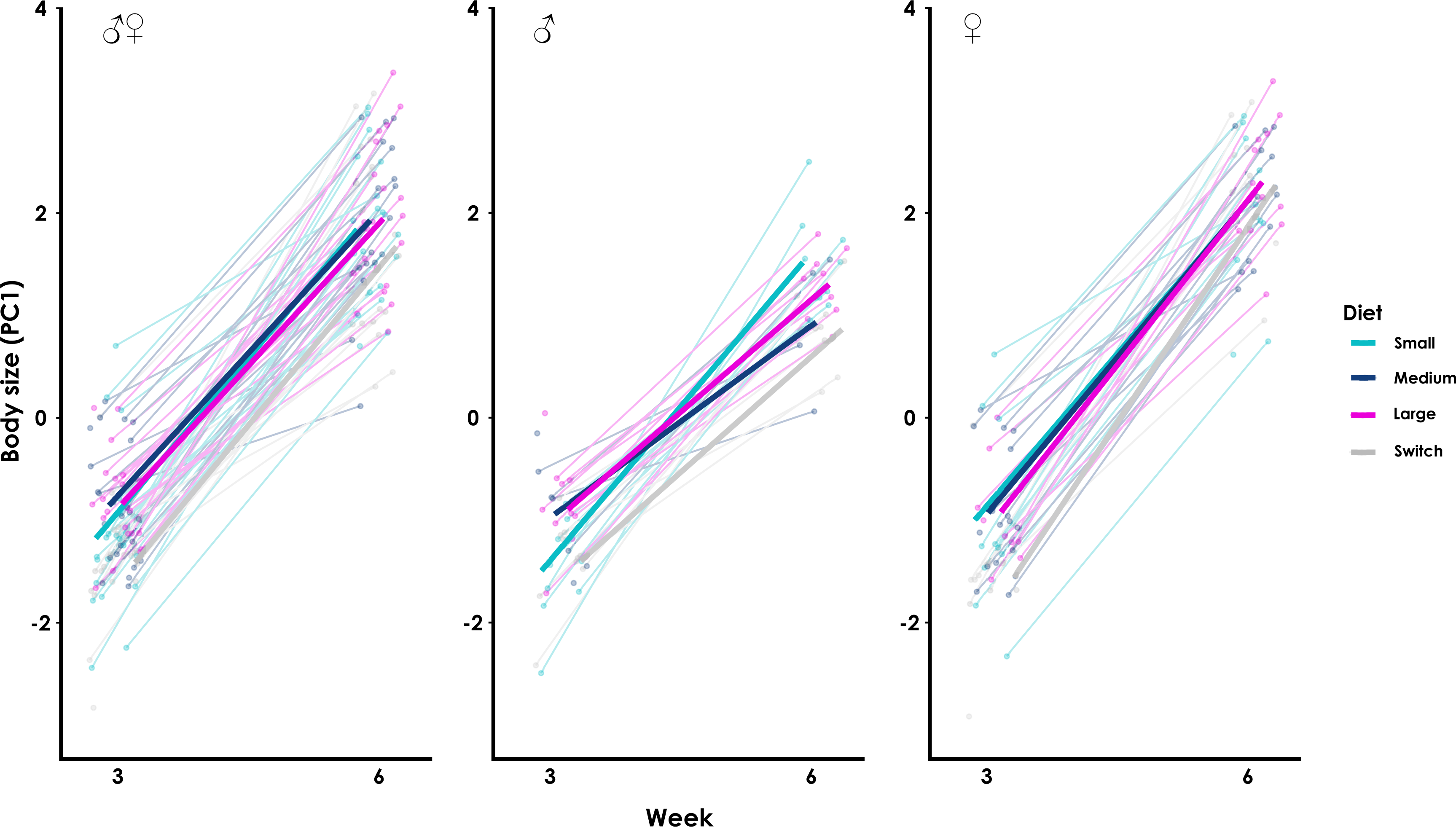
Lifetime change in body size (PC1) over age of Gryllodes sigillatus fed one of four different sizes of rabbit kibble: small (0.088-0.125 mm), medium (0.5-0.7 mm), large (1.0-1.4 mm), and a switch from small to large at 28 days after hatch. Thick lines connect mean values at each age and solid circles represent individual crickets connected by thin lines over time.

**Figure 4.**
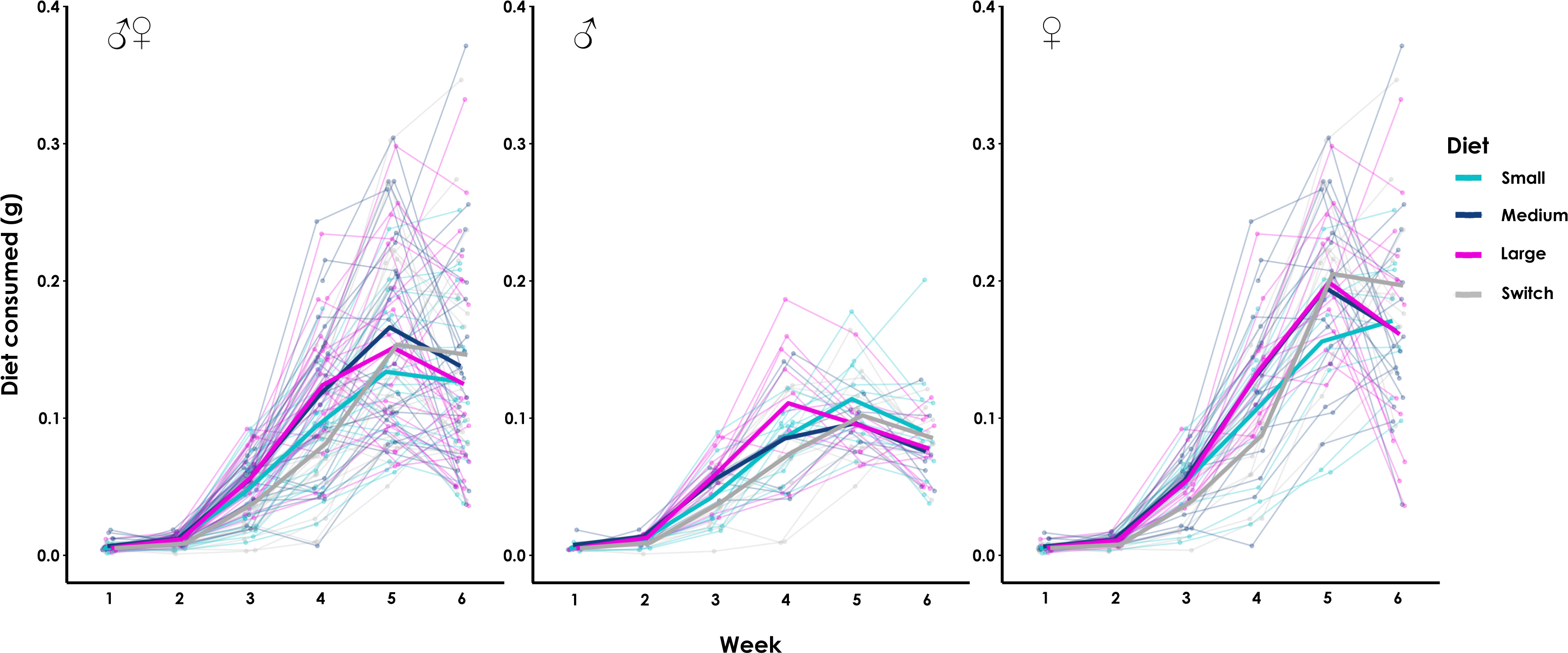
Lifetime consumption of food over age of Gryllodes sigillatus fed one of four different sizes of rabbit kibble: small (0.088-0.125 mm), medium (0.5-0.7 mm), large (1.0-1.4 mm), and a switch from small to large at 28 days after hatch. Thick lines connect mean values at each age and solid circles represent individual crickets connected by thin lines over time.

From the original 50 crickets used in the preference experiment, 21 females and 15 males survived the entire six weeks and were used in the data analysis. Of the 36 surviving crickets, all crickets reached adulthood except for 1 female and 1 male. Particle size had a significant influence on the amount of food consumed (F_2,464.15_ = 113.61; *P* < 0.001), and this effect changed over time (F_4,464_ = 119.98; *P* < 0.001). Crickets consumed significantly more of the medium and large diets compared to the small diet every week except for week two (Figure 5; Table S7). During weeks three, four, and six, crickets consumed significantly more large diet compared to the medium diet, and ate 12.5%, 46.2%, and 33.1% more food, respectively (Figure 5). During week five, crickets ate significantly more medium diet compared to the large diet and ate 40.0% more food (Figure 5).

**Figure 5.**
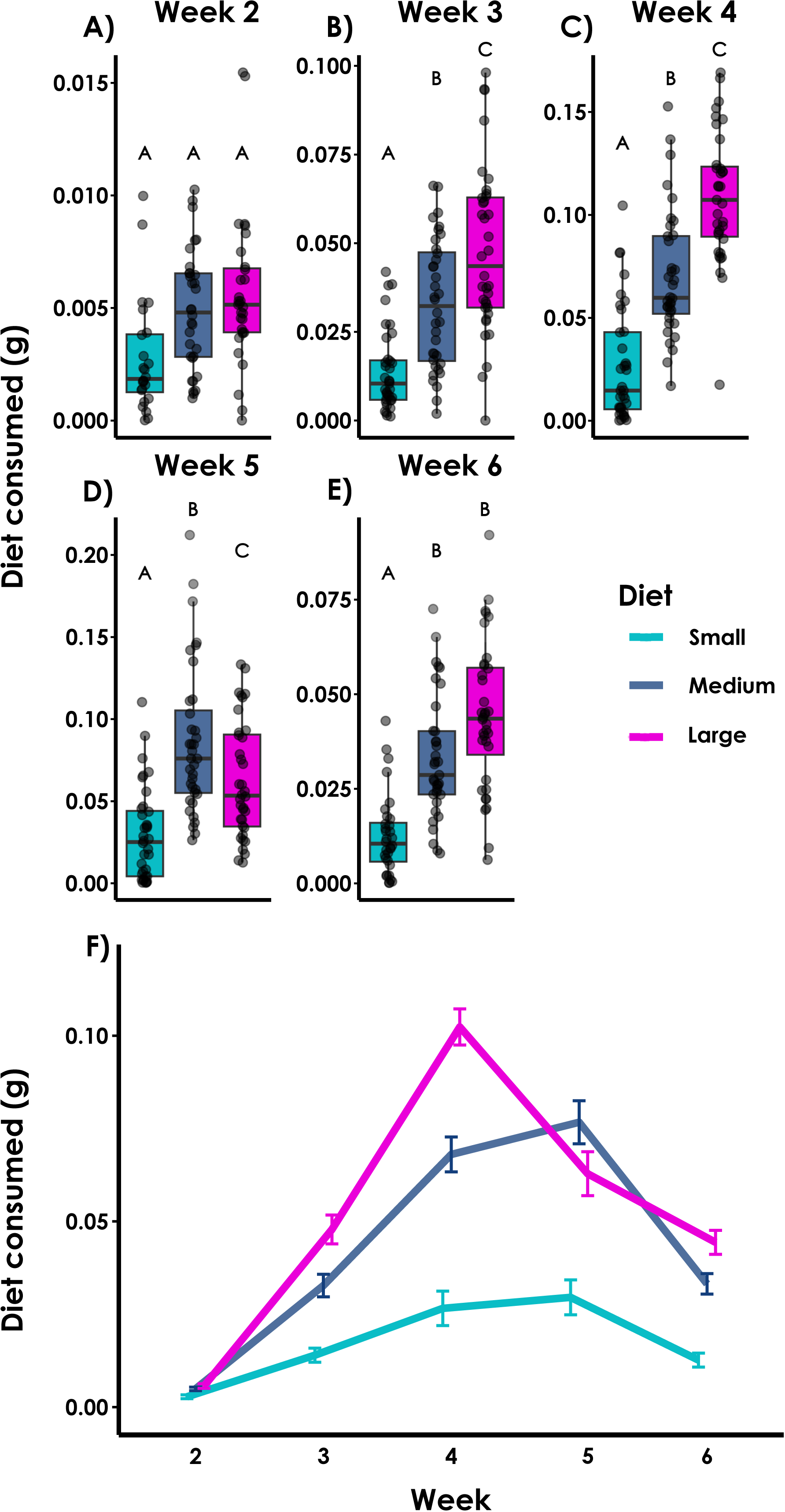
Weekly food consumption (preference) of individual Gryllodes sigillatus given the choice of three different sizes of rabbit kibble: small (0.088-0.125 mm), medium (0.5-0.7 mm), large (1.0-1.4 mm). A-E) The centre line for each box plot represents the median, and the bottom and top lines represent the 25th and 75th quantiles. The whiskers extend 1.5 times the interquartile range. Solid circles represent individual data points. Different letters between boxes within a week denote significant differences between diets. F) Thick lines connect mean values at each week, and error bars represent standard error.

### Pellet experiment

From the original 100 crickets, 18 females and 32 males survived for six weeks, reached adulthood, and were used in the data analysis. Pelleting the diet did not significantly influence cricket mass over time (F_1,116.32_ = 0.49; *P* = 0.49), and naturally crickets gained mass over time (F_5,326.72_ = 904.98; *P* < 0.001; Figure 6). Final mass at week five was also unaffected by diet (F_1,46_ = 2.54; *P* = 0.12) and there was no significant sex by diet interaction (F_1,46_ = 1.87; *P* = 0.18; Table S9; Figure 6). However, crickets fed the pelleted diet grew significantly larger than crickets fed the ground diet (Pellet: 0.68 ± 0.28 PC1; Control: 0.019 ± 0.26 PC1; F_1,46_ = 6.87; *P* = 0.012), and the interaction between sex and diet was not significant (F_1,46_ = 1.40; *P* = 0.24; Figure 6; Table S9).

**Figure 6.**
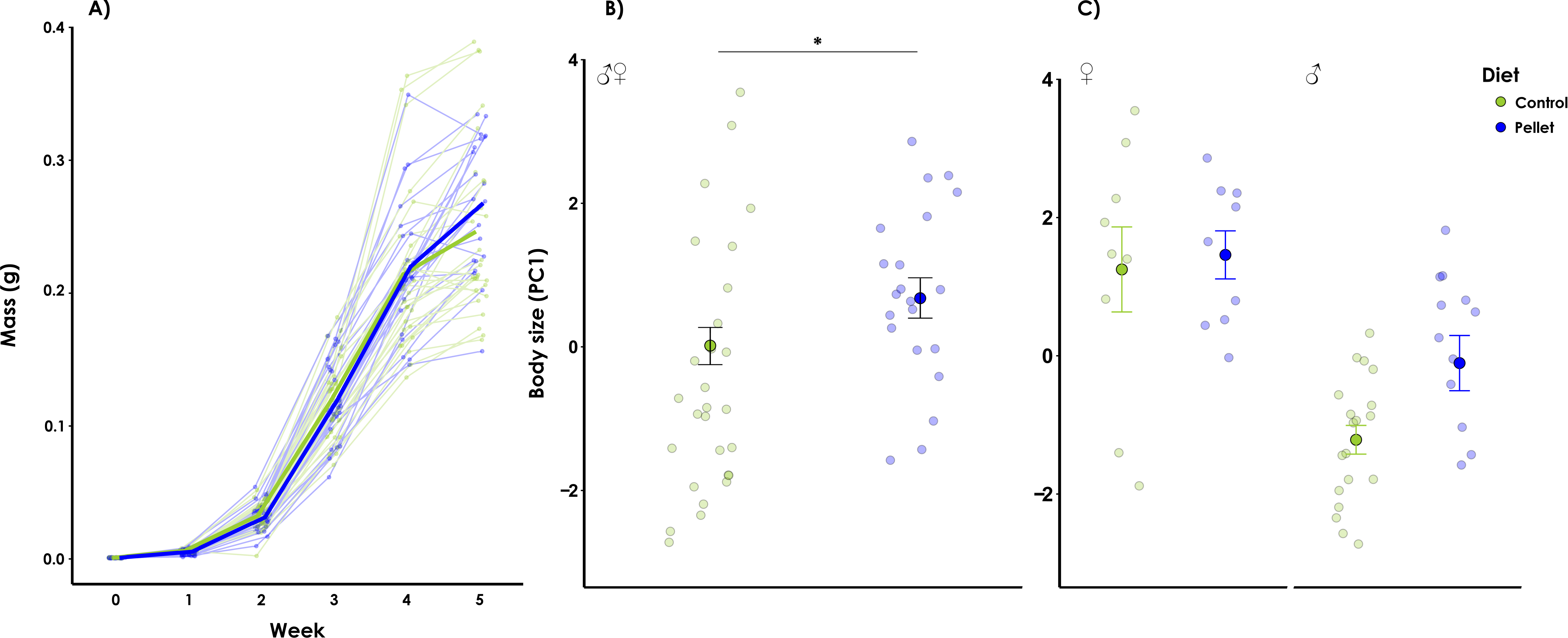
A) Weekly mass of crickets fed ground (control) and pelleted diets. Thick lines connect mean values at each age and solid circles represent individual crickets connected by thin lines over time. B) Adult body size (PC1) of pooled, C) male, and female crickets fed unpelleted (control) and pelleted diets. The pelleted diet was created by first grinding a conventional cricket diet, mixing it with tap water, and then drying and smearing over plexiglass molds to dry at 30°C. Transparent circles represent individual crickets, and darker larger circles represent least square means with standard error bars. Asterisk denotes significant difference (*P* < 0.05).

### Fine grinding experiments

There were no significant effects of grinding feed on any life history traits measured between the individual- and colony-level experiments (Figure 7). Fitted logistic growth curves for fine grind and untreated feed were similar to null models pooling feed type for individually-reared crickets (fine grind = 39, untreated = 40; likelihood ratio = 0.74, *P* = 0.95; Figure 7A) or for group-reared crickets (N = 20 per bin, 8 bins; likelihood ratio = 6.54, *P* = 0.16; Figure 7B). The effect of grinding feed on mean mass was not dependent on sex for 30 female and 27 male individually-reared crickets (F_1,196_ = 3.86; *P* = 0.051; Figures 7C,E) or between the 6^th^ and 8^th^ instars in group-reared (F_1,671_ = 0.36; *P* = 0.55; Figures 7D,F). The mean proportion of adults over time was similar between untreated and fine grind feed for group-reared (Z_69_ = -0.18; *P* = 0.85) and individually-reared crickets (Z_13_ = -0.36; *P* = 0.72). Biomass after seven weeks was similar between feed types for group-reared crickets (t_5.65_ = 0.43, *P* = 0.68; Figure 7H). Survival was also unaffected by feed treatment in group-reared crickets (Z_6_ = -0.07; *P* = 0.94; Figure 7J).

**Figure 7.**
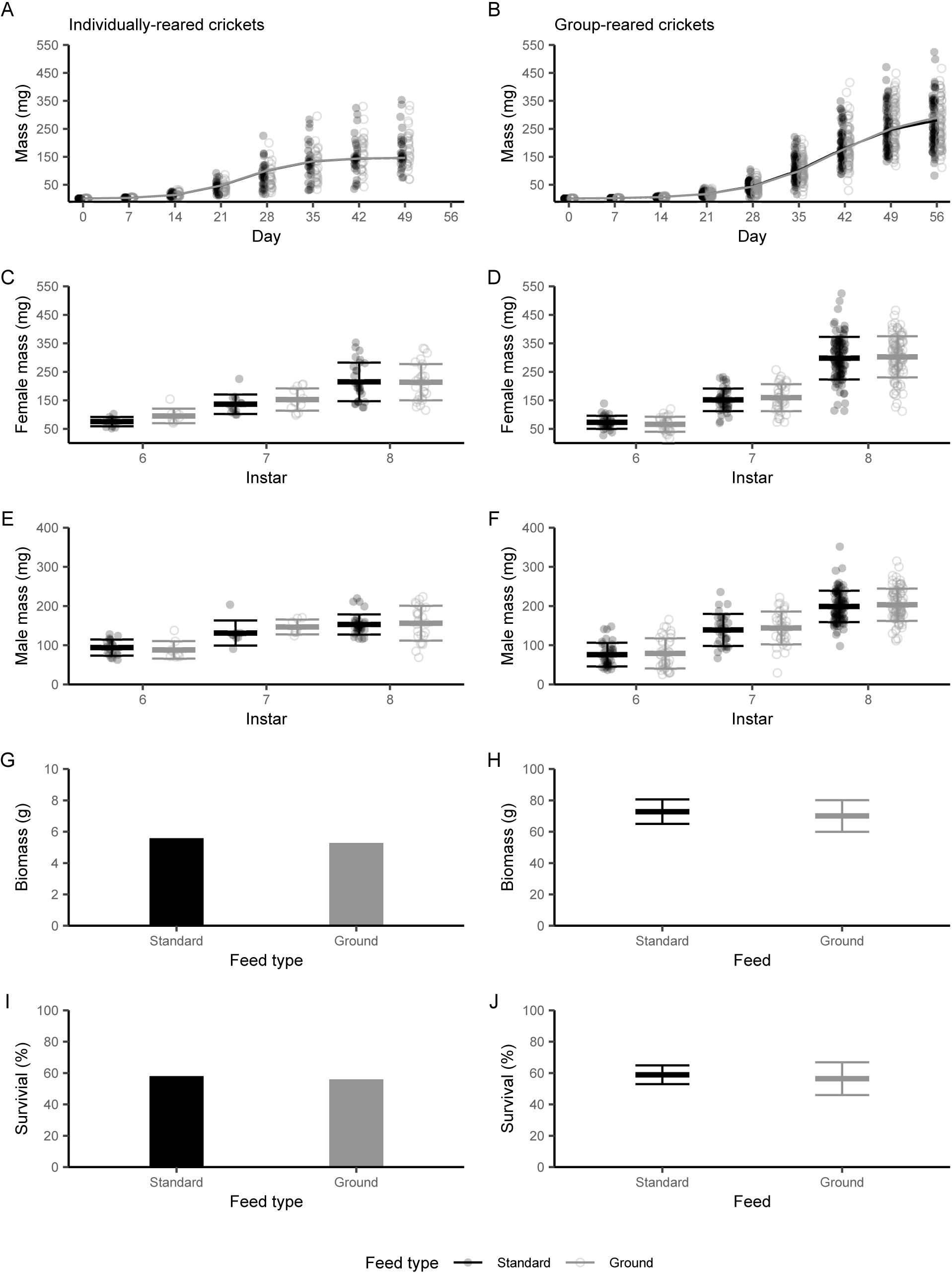
Life history traits of crickets fed unground commercial cricket feed (standard) and finely ground cricket feed (ground). Diet was ground using a stainless steel grain grinder on the finest possible setting, and both diets were fed to crickets from hatch until 7-weeks old. The left panel figures represent individually-reared crickets, and the right panel figures represent group-reared crickets. A,B) Pooled, C,D) female, and E,F) male mass of crickets fed both diets. Transparent circles represent individual crickets, thick bands represent the mean and thinner bands represent standard deviation. G,H) Total mass (biomass) of all remaining crickets after 7 weeks. I,J) Percent survival of crickets fed both diets.

## Discussion

Understanding the relationship between diet and life history traits can inform production optimization strategies. We ran a series of experiments that tested how grinding, sieving, and pelleting the diet influenced survival and growth of individual crickets, and whether crickets demonstrated a preference for a particular diet particle size. Here, we present novel results on the influence that dietary particle size can exert on cricket mass, body size, and development time: when restricted to one of three sized diets, crickets developed at a faster rate when fed the large diet (reaching adulthood approximately two days earlier) and grew heavier and larger bodies faster when fed medium and large sized diets compared to crickets fed small sized diet. When provided with a choice of size, crickets preferentially consumed more of the medium and large sized diets than the small sized diet. Importantly, our results represent an overall shift in the growth curve of *G. sigillatus*, and crickets developed into adults faster while consuming the same amount of food per week. This suggests that milling to an optimal particle size can contribute to scaling up production while reducing feed costs. Harvesting two days earlier could result in more product being produced in less time.

There is substantial potential for a larger sized diet to influence cricket mass production, especially considering most of the conventional cricket diet we ground and pelleted is smaller than the medium sized treatment diet. An interesting caveat is that under group-rearing conditions, *G. sigillatus* are observed to selectively feed on the conventional feed and leave behind larger sized particles (Figure S3), however this observation remains to be experimentally tested. It is unclear whether crickets are selectively feeding based on nutrient availability, particle size preference, taste, or a combination of these factors. Our experiments using rabbit kibble were specifically designed to prevent nutrient preference by using a homogenous diet. Future experiments should test the combined effects of particle and nutrient availability by designing homogenous and heterogenous isonutrient diets and then measuring food consumption and the relative amounts of remaining ingredients to ascertain which ingredients require more processing.

We attempted to restrict nutrient selection of the conventional cricket feed by pelleting the diet. The pellets we designed were ∼2x larger than the large diet used in the rabbit kibble no-choice experiment (2 mm vs 1.0-1.44 mm) and considering that large-sized rabbit kibble resulted in heavier and larger crickets earlier in development, it is surprising that large-sized pellets did not produce a similar result. Pelleting resulted in larger bodies, but not heavier mass, nor were there any significant effects throughout development. The pellets could have been too large; larger than the maximum particle size that optimizes cricket mass and body size. Our particle size experiments tested diets that ranged in size from 0.088 mm to 1.4 mm. However, our pellets were 2mm in diameter. It is possible, therefore, that *G. sigillatus* prefers a diet between 1.4 – 2.0 mm. More research is required that tests diets within this range to determine the full range of dietary size preference of *G. sigillatus* that results in a peak growth curve response. Also, testing a range of pellets that more closely resemble the peak particle size ranges would help to determine whether pelleting conventional cricket feed is an acceptable method for offering optimal particle sizes while restricting nutrient selection. A limitation of this study was that we did not test a ‘true’ pellet; pelletizing requires high pressure, temperature, and moisture achieved through steam (Shurson, 2018), but we tested the concept of a pellet with the resources and tools available to us and discovered that pelleting a cricket diet can modify cricket body size. Our particle size experiments suggest that homogenized diets can produce larger and bigger crickets when ground to an optimal particle size, and so future research should test diets that are formally pelletized to determine if homogenized and then pelleted feed can optimize cricket growth.

We found no support for our prediction that crickets fed the finely ground and smaller sized diet would grow the largest, which is surprising because reduction in particle size can increase production of other farmed animals (Goodband *et al*., 2002; Jo *et al*., 2021). However, there are records of negative effects from diets that are too small in particle size (ex. gastric ulcers in pigs; Ayles et al., 1996), and so the small particle size diet we tested (0.088-0.125 mm) may have been too small for *G. sigillatus* and thereby resulted in negative effects that counteracted any benefit. Regardless, crickets on the small diet still successfully developed into adults that were just as heavy as crickets fed the medium and large diets. The mechanisms behind the life history responses to and preference for larger diet particle size are not clear from our experiment, but we hypothesize that cricket hiding behaviour may influence selection for and performance on larger sized diet particles. Crickets hide to avoid predators, and common cricket rearing practice is to provide shelters for the crickets to hide within (Ayieko *et al*., 2016; Clifford *et al*., 1977; Hedrick and Kortet, 2006). Many insect species including crickets demonstrate cannibalism, where older and larger individuals prey on smaller juveniles, especially when the crickets are fed nutritionally-imbalanced diets (Gutiérrez *et al*., 2020; Richardson *et al*., 2010). *Acheta domesticus* has been observed handling food items one at a time (Tennis et al., 1979), and this may be true for other cricket species. Larger sized food particles may therefore enable crickets to grab-and-go: pick up a piece of food, bring it back to a hiding place, and spend more time eating while sheltered from predators. This idea is not, however, supported by our no-choice experiment on diet particle size as the amount of food consumed was unaffected by diet particle size.

However, in that experiment the rearing containers were small, with a piece of egg carton for shelter resting right above the food dish. As a result, the crickets could visit the feed dish and forage while simultaneously hiding. Although in the particle size choice experiment, crickets were held in larger rearing containers and had to venture out from hiding to feed and they ate more of the medium and large sized diets compared to the small diet. Future experiments could use cameras to monitor and measure how often individuals visit feed dishes relative to food particle size to determine whether crickets are transporting food away from the dish.

Our results provide clear evidence that researchers and insect producers should be thinking about and testing different particle sizes when designing insect diets. Feed is one of the most expensive components of cricket production (Morales-Ramos *et al*., 2020), and reduction in feed processing (i.e. less time and energy spent grinding feed) could reduce costs. Our results show that finely grinding conventional cricket feed does not influence mass, development time or survival. We also present evidence that crickets preferentially feed on a large sized diet (1.0 – 1.4 mm) throughout development and were more likely to develop faster than crickets fed a small sized diet (0.088 – 0.125 mm). We also demonstrated that pelleting cricket diet has potential to manipulate body size, but more research testing formally pelleted insect diets is required before implementing these findings in a large-scale setting.

## Acknowledgements

We would like to thank Dr. Jeff Dawson for his instruction and lending of his drill press to create pellet molds. Thank you to Celine Larose for ensuring we always had a healthy supply of pelleted diet. This work was supported by a MITACS Accelerate Internship, funding from the Province of Ontario, Ontario Graduate Scholarships awarded to M.J.M., NSERC Discovery Grants awarded to both H.A.M. and S.M.B., and the Canadian Foundation for Innovation.

