## Supplemental tables and figures for "Larger diet particle sizes cause crickets to grow faster with no effect on final body size"

**Table S1.** Ingredient list for the rabbit kibble (Martin Little Friends, Martin Mills Inc., Elmira, Ontario, Canada). Ingredients are copied directly from the product packaging.

| **Ingredient** |
| --- |
| Ascorbic acid |
| Barley |
| Biotin |
| Brewers yeast |
| Calcium carbonate |
| Calcium iodate |
| Calcium pantothenate |
| Choline chloride |
| Cobalt sulphate |
| Copper sulphate |
| Dehydrated alfalfa |
| Dicalcium phosphate |
| DL-methionine |
| Folic acid |
| Iron sulphate |
| Lignin sulfonate |
| Manganese sulfate |
| Mannan oligosaccharide |
| Mineral oil |
| Niacinamide |
| Pyridoxine hydrochloride |
| Riboflavin |
| Rice hulls |
| Salt |
| Selenium |
| Soya oil (preserved with rosemary extract, mixed tocopherols, and citric acid) |
| Soybean hulls |
| Soybeans |
| Thiamine mononitrate |
| Vitamin A |
| Vitamin B12 |
| Vitamin D3 |
| Vitamin E |
| Vitamin K |
| Wheat shorts |
| Yucca schidigera extract |
| Zinc oxide |

**Table S2.** Guaranteed proximate analysis of rabbit kibble (Martin Little Friends, Martin Mills Inc., Elmira, Ontario, Canada). Values are copied directly from the product packaging.

| Ingredient | Percent composition |
|---|---:|
| Crude protein | 16% |
| Crude fat | 4% |
| Crude fibre | 15-18% |
| Moisture | 10% |
| Sodium | 0.2% |
| Calcium | 0.9% |
| Phosphorus | 0.5% |
| Vitramin A | 17’250 I.U./kg |
| Vitamin D3 | 2’150 I.U./kg |
| Vitamin E | 54 I.U./kg |

**Table S3.** Model results from Type II analyses of variance associated with the linear models of how, when not given a choice, diet particle size influenced mass and body size (PC1) of individual crickets. Models were performed on pooled sex data at week 3 and 6, and then also separately on males and females at week 6. The week 3 models pooled the small and switch diet had not yet changed from small to large particle size. These diets were not pooled for the week 6 models, as the switch diet had changed from small to large. Detailed linear model results found in Table S3.

| **Model** | **Week** | **Term** | ***df*** | ***F*** | ***P*** |
|---|---|---|---|---|---|
| Mass | 3 | **Diet** | **2, 86** | **11.09** | **< 0.001** |
| | | Sex | 1, 86 | 0.10 | 0.75 |
| | | Diet*Sex | 2, 86 | 0.20 | 0.82 |
| | 6 | Diet | 3, 81 | 1.14 | 0.34 |
| | | **Sex** | **1, 81** | **53.43** | **< 0.001** |
| | | Diet*Sex | 3, 81 | 0.36 | 0.78 |
| Body size (PC1) | 3 | **Diet** | **2, 84** | **10.11** | **< 0.001** |
| | | Sex | 1, 84 | 0.075 | 0.78 |
| | | Diet*Sex | 2, 84 | 0.0075 | 0.99 |
| | 6 | Diet | 3, 69 | 0.73 | 0.54 |
| | | **Sex** | **1, 69** | **50.04** | **< 0.001** |
| | | Diet*Sex | 3, 69 | 1.86 | 0.15 |

**Table S4.** The model results from the linear models of how, when not given a choice, diet particle size influenced mass and body size (PC1) of individual crickets at week 3 and week 6. The intercept for week 3 corresponds to diet = large and sex = male, and the intercept for week 6 corresponds to diet = small and sex = male.

| **Model** | **Week** | **R2** | **Term** | **Estimate (± SE)** | **t value** | ***P*** |
|---|---|---|---|---|---|---|
| Mass | 3 | 0.16 | Intercept | 0.076 ± 0.0059 | 12.83 | < 0.001 |
| | | | Medium | -0.0033 ± 0.0095 | -0.35 | 0.73 |
| | | | **Small** | **-0.023 ± 0.0073** | **-3.17** | **< 0.01** |
| | | | Female | -0.00087 ± 0.0082 | -0.11 | 0.92 |
| | | | Medium*Female | -0.0053 ± 0.012 | -0.44 | 0.66 |
| | | | Small*Female | 0.0014 ± 0.010 | 0.14 | 0.89 |
| | 6 | 0.37 | **Intercept** | **0.19 ± 0.013** | **14.29** | **< 0.001** |
| | | | Medium | -0.0072 ± 0.021 | -0.34 | 0.73 |
| | | | Large | -0.00092 ± 0.019 | -0.049 | 0.96 |
| | | | Switch | -0.019 ± 0.019 | -1.00 | 0.32 |
| | | | **Female** | **0.059 ± 0.020** | **3.00** | **< 0.01** |
| | | | Medium*Female | 0.016 ± 0.028 | 0.59 | 0.56 |
| | | | Large*Female | 0.0028 ± 0.026 | 0.11 | 0.92 |
| | | | Switch*Female | 0.024 ± 0.027 | 0.89 | 0.38 |
| Body size (PC1) | 3 | 0.15 | **Intercept** | **-1.13 ± 0.20** | **-5.66** | **< 0.001** |
| | | | Medium | -0.047 ± 0.32 | -0.15 | 0.89 |
| | | | **Small** | **-0.66 ± 0.25** | **-2.67** | **< 0.01** |
| | | | Female | 0.0096 ± 0.28 | 0.034 | 0.97 |
| | | | Medium*Female | 0.039 ± 0.41 | 0.095 | 0.93 |
| | | | Small*Female | 0.041 ± 0.35 | 0.12 | 0.91 |
| | 6 | 0.40 | **Intercept** | **1.40 ± 0.19** | **7.25** | **< 0.001** |
| | | | Medium | -0.41 ± 0.35 | -1.17 | 0.25 |
| | | | Large | -0.22 ± 0.27 | -0.82 | 0.42 |
| | | | **Switch** | **-0.55 ± 0.27** | **-2.04** | **0.046** |
| | | | Female | 0.48 ± 0.26 | 1.86 | 0.068 |
| | | | Medium*Female | 0.57 ± 0.42 | 1.36 | 0.18 |
| | | | Large*Female | 0.56 ± 0.36 | 1.54 | 0.13 |
| | | | **Switch*Female** | **0.86 ± 0.37** | **2.30** | **0.024** |

**Table S5.** The model results from the linear mixed model of how, when not given a choice, diet particle size influenced food consumption of individual crickets. Cricket ID was included as a random effect. The intercept corresponds to Week = 0, Diet = Small, and Sex = Male.

| Model | R2 | Term | Estimate (± SE) | df | t value | *P* |
|---|---|---|---|---|---|---|
| Food consumption | 0.66 | **Intercept** | **-0.018 ± 0.0064** | **229.56** | **-2.74** | **0.0066** |
| | | Week 2 | 0.0056 ± 0.0062 | 450.19 | 0.91 | 0.36 |
| | | **Week 3** | **0.045 ± 0.0062** | **450.79** | **7.26** | **< 0.001** |
| | | **Week 4** | **0.099 ± 0.0062** | **450.79** | **16.07** | **< 0.001** |
| | | **Week 5** | **0.15 ± 0.0062** | **452.89** | **23.67** | **< 0.001** |
| | | **Week 6** | **0.13 ± 0.0062** | **452.95** | **20.75** | **< 0.001** |
| | | Medium | 0.0056 ± 0.0062 | 90.75 | 0.90 | 0.37 |
| | | Large | 0.0077 ± 0.0061 | 90.54 | 1.25 | 0.22 |
| | | Switch | 0.0014 ± 0.0062 | 90.84 | 0.23 | 0.82 |
| | | **Female** | **0.033 ± 0.0044** | **89.68** | **7.58** | **< 0.001** |

**Table S13.** Least squares means ± standard error of the amount of food consumed (g) of individual crickets from the no-choice diet particle size experiment. Values within a column and term followed by different letters are significantly different ($P < 0.05$).

| Term | Level | Food consumed (g) ± SE | |
|---|---|---|---|
| Diet | Small | 0.070 ± 0.0045 | A |
| | Switch | 0.071 ± 0.0043 | A |
| | Medium | 0.076 ± 0.0043 | A |
| | Large | 0.078 ± 0.0042 | A |
| Week | Week 1 | 0.0028 ± 0.0046 | A |
| | Week 2 | 0.0084 ± 0.0045 | A |
| | Week 3 | 0.048 ± 0.0045 | B |
| | Week 4 | 0.10 ± 0.0045 | C |
| | Week 5 | 0.15 ± 0.0046 | D |
| | Week 6 | 0.13 ± 0.0046 | E |
| Sex | Male | 0.057 ± 0.0033 | A |
| | Female | 0.090 ± 0.0029 | B |

**Table S14.** Model results from linear mixed model of how diet particle size influenced food consumption of individual crickets when given the choice of three different sizes of rabbit kibble: small (0.088-0.125 mm), medium (0.5-0.7 mm), large (1.0-1.4 mm). Cricket ID was included as a random effect. The intercept corresponds to Week = 2 and Diet = Small.

| **Model** | **R2** | **Term** | **Estimate (± SE)** | **df** | **t-value** | ***P*** |
|---|---|---|---|---|---|---|
| Food consumed | 0.60 | Intercept | 0.0023 ± 0.0050 | 473.4 | 0.46 | 0.65 |
| | | Week 3 | 0.012 ± 0.0062 | 470.5 | 1.93 | 0.054 |
| | | **Week 4** | **0.024 ± 0.0062** | **470.1** | **3.98** | **< 0.001** |
| | | **Week 5** | **0.027 ± 0.0062** | **470.1** | **4.46** | **< 0.001** |
| | | Week 6 | 0.011 ± 0.0063 | 471.8 | 1.75 | 0.081 |
| | | Medium | 0.0024 ± 0.0063 | 470.3 | 0.38 | 0.71 |
| | | Large | 0.0030 ± 0.0063 | 473 | 0.47 | 0.64 |
| | | Week 3*Medium | 0.016 ± 0.0083 | 466.9 | 1.95 | 0.052 |
| | | **Week 4*Medium** | **0.041 ± 0.0083** | **466.7** | **5.00** | **< 0.001** |
| | | **Week 5*Medium** | **0.054 ± 0.0083** | **466.7** | **6.53** | **< 0.001** |
| | | **Week 6*Medium** | **0.018 ± 0.0084** | **467.8** | **2.10** | **0.036** |
| | | **Week 3*Large** | **0.031 ± 0.0084** | **468.6** | **3.68** | **< 0.001** |
| | | **Week 4*Large** | **0.079 ± 0.0084** | **468.1** | **9.50** | **< 0.001** |
| | | **Week 5*Large** | **0.030 ± 0.0083** | **468.4** | **3.64** | **< 0.001** |
| | | **Week 6*Large** | **0.028 ± 0.0085** | **469.2** | **3.33** | **< 0.001** |

**Table S8.** Model result from a Type III analysis of variance associated with the linear mixed model of how diet particle size influenced food consumption of individual crickets when given the choice of three different sizes of rabbit kibble: small (0.088-0.125 mm), medium (0.5-0.7 mm), large (1.0-1.4 mm). Cricket ID was included as a random effect.

| **Model** | **Term** | ***df*** | ***F*** | ***P*** |
|---|---|---|---|---|
| Food consumed | **Week** | **4, 464** | **119.98** | **< 0.001** |
| | **Diet** | **2, 464.15** | **113.61** | **< 0.001** |
| | **Week*Diet** | **8, 463.76** | **18.45** | **< 0.001** |

**Table S12.** Model results from Type III analyses of variance associated with the linear models of how pelleting a conventional cricket diet influenced the mass and body size of individual crickets.

| Model | Term | *df* | *F* | *P* |
|---|---|---|---|---|
| Mass | Diet | 1, 46 | 2.54 | 0.12 |
| | **Sex** | **1, 46** | **27.47** | **< 0.001** |
| | Diet*Sex | 1, 46 | 1.87 | 0.18 |
| Body size (PC1) | **Diet** | **1,46** | **6.87** | **0.012** |
| | **Sex** | **1,46** | **29.07** | **< 0.001** |
| | Diet*Sex | 1,46 | 1.40 | 0.24 |

**Table S13.** SIZE NO CHOICE - Penultimate confidence set of linear mixed models examining which predictors (Diet, Week, and Sex) and their interaction best explained the amount of diet consumed.

| Variable | Model | AIC | Δ AIC [a] | Weight [b] | Evidence Ratio [c] | Model selected for analysis |
|---|---|---|---|---|---|---|
| Diet consumed | Diet*Week | -1636.92 | 145.95 | 2.03e-32 | 1.00E | No |
| | Diet + Week | -1748.90 | 34.00 | 4.20e-08 | 2.07e+24 | No |
| | **Diet + Week + Sex** | **-1782.87** | **0.00** | **1.00** | **4.93e+31** | **Yes** |
| | Diet*Week*Sex | -1640.19 | 143.00 | 1.04e-31 | 5.14 | No |
| | Diet*Week + Sex | -1670.66 | 112.00 | 4.30e-25 | 2.12e+07 | No |

[a] The difference between the current model's AIC score and that of the top ranked model (bolded).

[b] Akaike weights, or relative likelihood of the current model. Calculated as exp(-0.5 x Δ AIC) divided by the sum of these values across all models in the confidence set.

[c] Evidence ratio, or how many times more parsimonious the top ranked model (bolded) is over the current model. Calculated as the Akaike weight of the top ranked model divided by that of the current model.

**Table S14. SIZE – CHOICE -** Penultimate confidence set of linear mixed models examining which predictors (Diet, Week, and Sex) and their interaction best explained amount of diet consumed.

| Variable | Model | AIC | Δ AIC [a] | Weight [b] | Evidence Ratio [c] | Model selected for analysis |
|---|---|---|---|---|---|---|
| Diet consumed | **Diet*Week** | **-2213.50** | **0.00** | **9.28e-01** | **1.37e+17** | **Yes** |
| | Diet + Week | -2165.36 | 48.14 | 3.26e-11 | 4.81e+06 | No |
| | Diet*Week*Sex | -2134.59 | 78.92 | 6.78e-18 | 1.00 | No |
| | Diet*Week + Sex | -2208.39 | 5.12 | 7.18e-02 | 1.06e+16 | No |
| | Diet + Week + Sex | -2163.39 | 50.11 | 1.22e-11 | 1.80e+06 | No |

[a] The difference between the current model's AIC score and that of the top ranked model (bolded). Models with AIC > 4 have been omitted from this table

[b] Akaike weights, or relative likelihood of the current model. Calculated as exp(-0.5 x Δ AIC) divided by the sum of these values across all models in the confidence set.

[c] Evidence ratio, or how many times more parsimonious the top ranked model (bolded) is over the current model. Calculated as the Akaike weight of the top ranked model divided by that of the current model.

**Table S12.** PELLET **-** Penultimate confidence set of linear mixed models examining which predictors (Diet, Week, and Sex) and their interaction best explained amount of food consumed.

| Variable | Model | AIC | Δ AIC [a] | Weight [b] | Evidence Ratio [c] | Model selected for analysis |
|---|---|---|---|---|---|---|
| Food consumed | Diet*Week | -1474.60 | 41.94 | 7.83e-10 | 4.23e+91 | No |
| | **Diet + Week** | **-1516.54** | **0.00** | **1.00** | **5.40e+100** | **Yes** |
| | Diet + Week + Sex | -1105.03 | 411.51 | 4.38e-90 | 2.37e+11 | No |
| | Diet*Week + Sex | -1063.52 | 453.02 | 4.26e-99 | 2.30e+02 | No |
| | Diet*Week*Sex | -1052.65 | 463.89 | 1.85e-101 | 1.00 | No |

[a] The difference between the current model's AIC score and that of the top ranked model (bolded).

[b] Akaike weights, or relative likelihood of the current model. Calculated as exp(-0.5 x Δ AIC) divided by the sum of these values across all models in the confidence set.

[c] Evidence ratio, or how many times more parsimonious the top ranked model (bolded) is over the current model. Calculated as the Akaike weight of the top ranked model divided by that of the current model.

**Table S13.** The model results from the linear mixed model of how pelleting a conventional cricket diet (pelleted) influenced mass of individual crickets. The intercept corresponds to Week = 0 and Diet = control.

| Model | R2 | Term | Estimate (± SE) | df | t value | *P* |
|---|---|---|---|---|---|---|
| Mass | 0.91 | Intercept | -0.00051 ± 0.0038 | 251.1 | -0.14 | 0.89 |
| | | Week 1 | 0.0049 ± 0.0042 | 327.3 | 1.16 | 0.25 |
| | | **Week 2** | **0.031 ± 0.0049** | **343.8** | **6.38** | **< 0.001** |
| | | **Week 3** | **0.12 ± 0.0049** | **344.2** | **24.00** | **< 0.001** |
| | | **Week 4** | **0.22 ± 0.0049** | **344.2** | **43.76** | **< 0.001** |
| | | **Week 5** | **0.26 ± 0.0049** | **344.2** | **51.66** | **< 0.001** |
| | | Pelleted | 0.0029 ± 0.0042 | 116.3 | 0.70 | 0.487 |

**Table S14.** Model results from the linear model of how pelleting a conventional cricket diet (pelleted) influenced body size (PC1) of individual crickets. The intercept corresponds to Diet = control and Sex = Female.

| R2 | Term | Estimate (± SE) | t value | *P* |
|---|---|---|---|---|
| 0.41 | **Intercept** | **1.25 ± 0.43** | **2.93** | **< 0.01** |
| | Pelleted | 0.21 ± 0.61 | 0.35 | 0.73 |
| | **Male** | **-2.46 ± 0.52** | **-4.75** | **< 0.001** |
| | Pelleted*Male | 0.90 ± 0.76 | 1.18 | 0.24 |

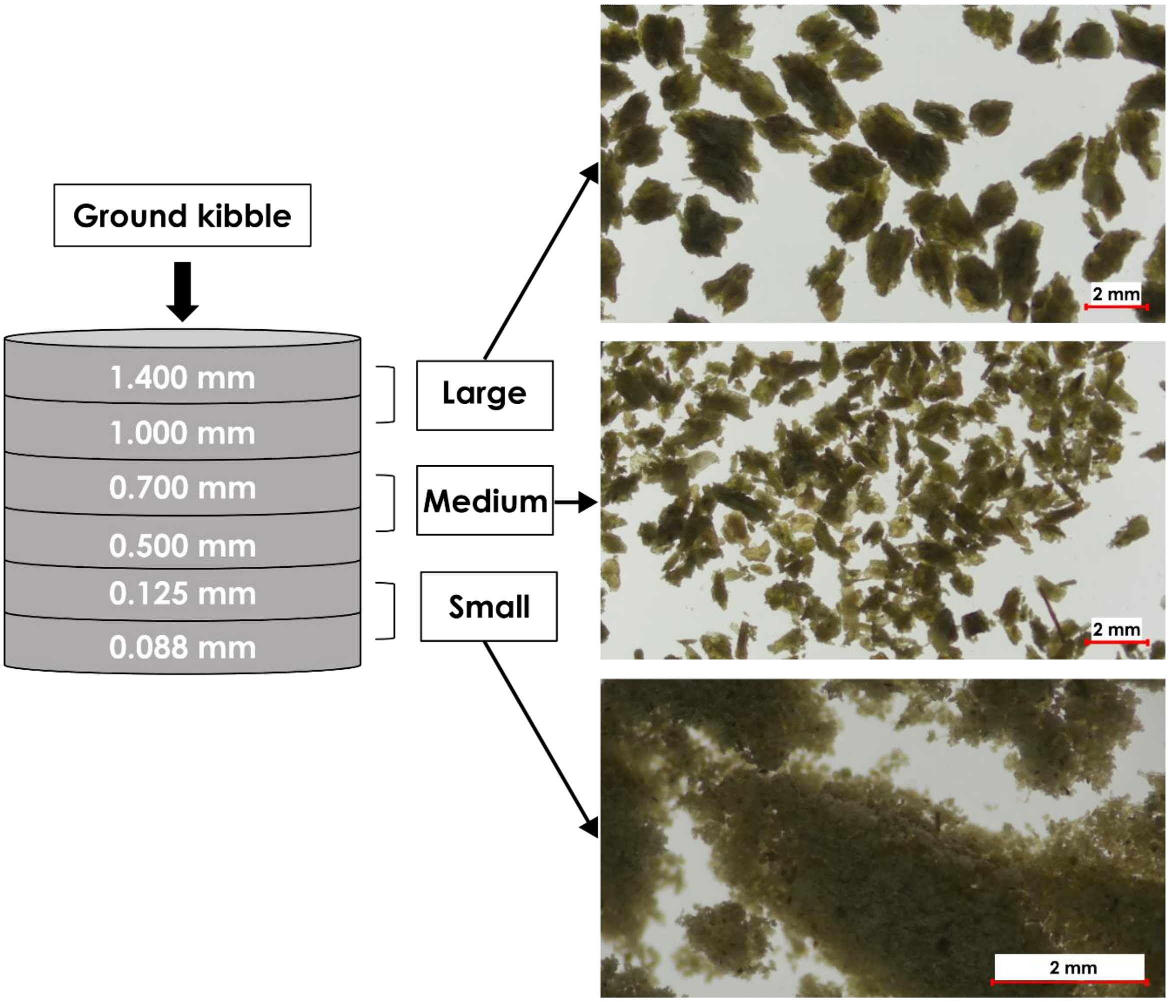


**Figure S1.** Sieve series used to separate ground rabbit kibble by particle size.

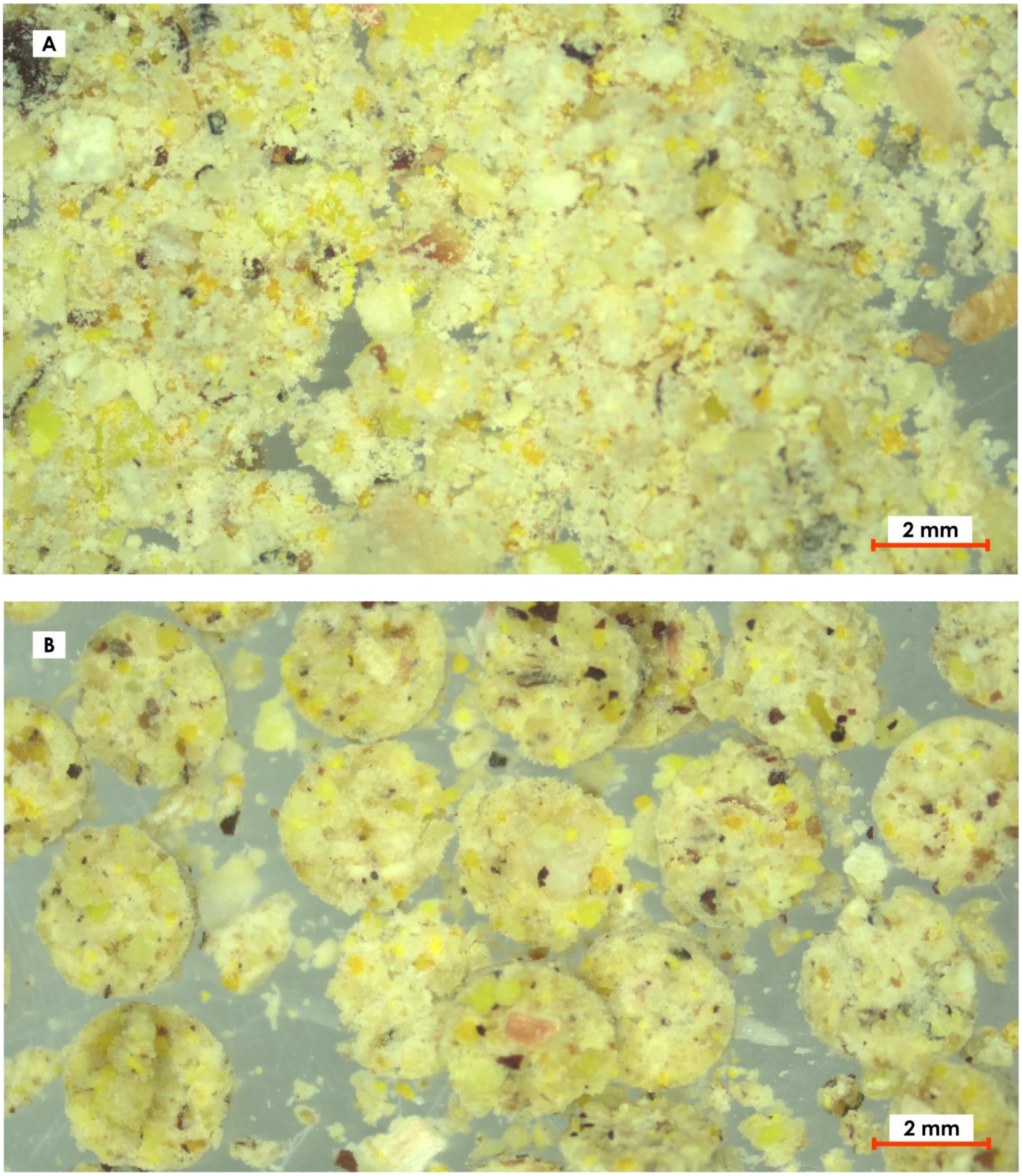


**Figure S2.** Ground conventional cricket feed prior to pelleting (A) and pelleted cricket feed (B).

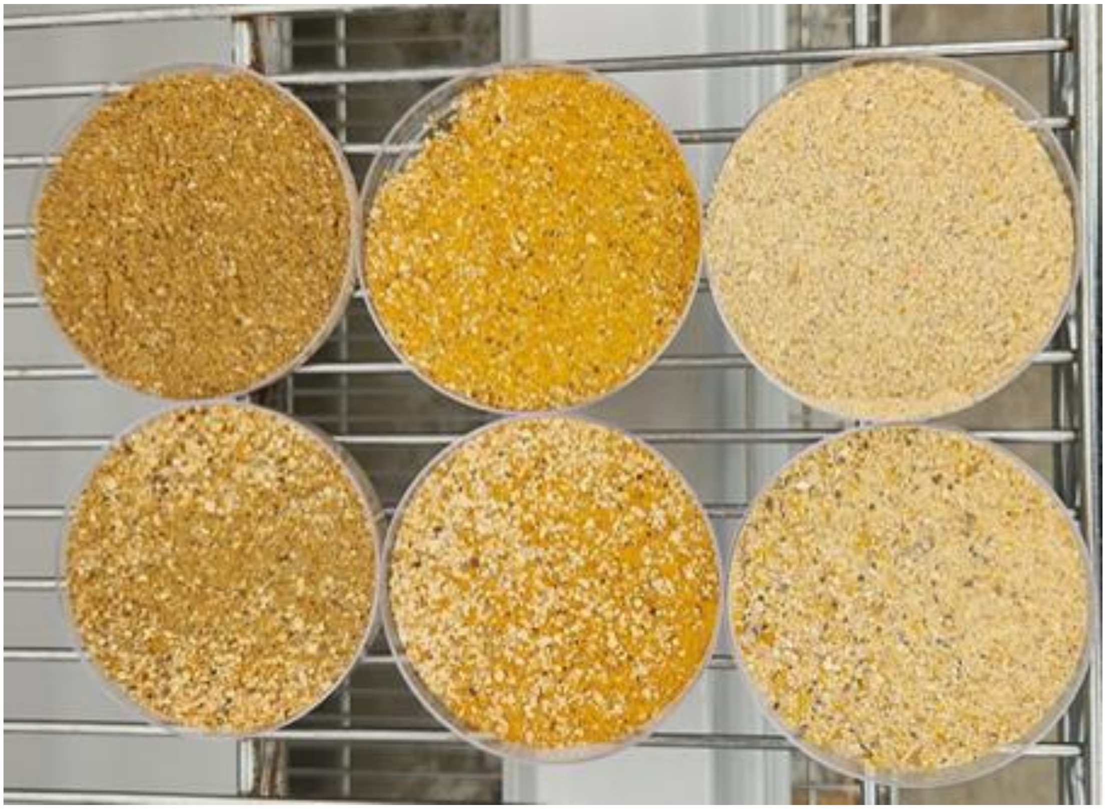

**Figure S3.** Feed dishes demonstrating selective feeding by group reared *Gryllodes sigillatus*. Dishes in the top row contain fresh diet that has not yet been fed upon. Dishes in the bottom row contain feed that has been fed upon for one week. The two dishes on the far right contain the conventional cricket feed used for the particle size and pellet experiments, whereas the four other dishes contain modifications of the conventional cricket feed for use in separate experiments.
